# SARS-CoV2 spike protein displays biologically significant similarities with paramyxovirus surface proteins; a bioinformatics study

**DOI:** 10.1101/2020.07.20.210534

**Authors:** Ehsan Ahmadi, Mohammad Reza Zabihi, Ramin Hosseinzadeh, Farshid Noorbakhsh

**Author notes:** **Correspondence to:** Dr Farshid Noorbakhsh, Department of Immunology, School of Medicine, Tehran University of Medical Sciences, Tehran, Iran.

## Abstract

Recent emergence of SARS-CoV2 and associated COVID-19 pandemic has posed a great challenge for the scientific community. Understanding various aspects of SARS-CoV2 biology, virulence and pathogenesis as well as determinants of immune response have become a global research priority. In this study, we performed bioinformatic analyses on SAR-CoV2 protein sequences, trying to unravel biologically important similarities between this newly emerged virus with other RNA viruses. Comparing the proteome of SARS-CoV2 with major positive and negative strand ssRNA viruses showed significant homologies between SARS-CoV2 spike protein with pathogenic paramyxovirus fusion proteins. This ‘spike-fusion’ homology was not limited to SARS-CoV2 and it existed for some other pathogenic coronaviruses; nonetheless, SARS-CoV2 spike-fusion homology was orders of magnitude stronger than homologies observed for other known coronaviruses. Moreover, this homology did not seem to be a consequence of general ssRNA virus phylogenetic relations. We also explored potential immunological significance of this spike-fusion homology. Spike protein epitope analysis using experimentally verified data deposited in Immune Epitope Database (IEDB) revealed that the majority of spike’s T cell epitopes as well as many B cell and MHC binding epitopes map within the spike-fusion homology region. Overall, our data indicate that there might be a relation between SARS-CoV2 and paramyxoviruses at the level of their surface proteins and this relation could be of crucial immunological importance.

## Introduction

Current COVID-19 pandemic which is caused by a newly discovered betacoronavirus has led to immense health and socio-economic problems around the world. First human coronaviruses were discovered in 1960s, but human coronavirus infections have likely existed for millennia (1–3). In the first few decades after their discovery, and when compared with other major viral pathogens like smallpox, polio and influenza, coronaviruses seemed to display a more benign attitude towards their hosts. Nonetheless, emergence of three deadly coronavirus epidemics over the last 20 years has challenged this view and is likely indicative of a new dynamism in the evolution of these pathogens (4, 5). This necessitates investigations that can provide more insight regarding coronavirus genes, proteins and biological behavior. Complex relations exist between viral pathogens at various evolutionary, epidemiological and pathogenic levels. One approach towards better understanding of a new pathogen is comparative analysis of its nucleic acids, proteins and molecular structures. The results of these types of analyses can go beyond ordinary phylogenetics and molecular biology and might have applied clinicopathological implications.

SARS-CoV2 nucleic acid and protein sequences did become available shortly after the start of COVID-19 pandemic and biological aspects of these genes/proteins including their interactions with immune system are under intense investigation. Recently, several reports have suggested that immunization against other human pathogens might influence susceptibility to SARS-CoV2. Live bacterial and viral vaccines including BCG, polio and MMR have been suggested as candidates that might give rise to this potential ‘cross-protective’ phenomenon (6–9). Considering the phylogenetic disparity between mycobacteria and RNA viruses, the effects of BCG have been mostly attributed to alterations at the level of innate immunity; i.e. immune-training (9). However, beneficial effects of unrelated viral vaccines might be due to innate or adaptive immune mechanisms, the latter being a consequence of similar antigenic structures.

In this study, we performed a comprehensive bioinformatic analysis on SARS-CoV2 protein sequences to determine whether there might be any biologically significant similarities between SARS-CoV2 proteins with non-coronavirus ssRNA viruses. We performed protein homology searches for individual SARS-CoV2 proteins against protein sequences from all major ssRNA virus families. This reveled some interesting similarities between SARS-CoV2 spike protein and paramyxovirus surface proteins as well as a SARS-CoV2 non-structural protein with Togaviruses. Considering the importance of spike proteins in viral infectivity and immune response, we then performed more extensive analyses on other coronavirus spike proteins as well as analyses to determine potential immunological significance of these homologies.

## Methods

### Extracting SARS-CoV2 and other coronavirus protein sequences

SARS-CoV2 Reference Protein sequences were obtained from NCBI Protein database (https://www.ncbi.nlm.nih.gov/protein). SARS-CoV2 RefSeq accession numbers and their associated official gene games are shown in Supplemental Table 1. Sequences for the surface proteins of various alpha, beta, gamma and delta-coronaviruses were also obtained from NCBI Protein database (Supplemental Table 2).

### Protein homology searches

Protein sequence homology searches were performed for all SARS-CoV2 proteins against major positive and negative strand ssRNA virus families using BLASP and Delta-BLAST algorithms. To detect and report the strongest homologies, best E value-alignment score pairs were extracted for each query. Likewise, protein homology searches were performed for spike proteins of alpha, beta, delta and gamma coronaviruses against ssRNA virus families. Results were shown as negative log10 of E values.

### Determining SARS-CoV2 spike protein epitopes and their distribution

Peptide sequences from SARS-CoV2 spike protein which could act as B cell or T cell epitopes or MHC binders were determined using immune epitope database (IEDB). Experimentally verified epitopes were obtained by searching the IEDB database with SARS-CoV2 spike protein. Linear epitopes were considered that had exact matches with substrings of SARS-CoV2 spike protein sequence. Pairwise BLASTP search was used to find the distribution of these epitopes on spike protein.

### Phylogenetic analysis

Phylogenetic analysis was performed using RNA-dependent RNA polymerase (RdRP) protein sequences from different ssRNA viruses. RdRP sequences from representative positive strand and negative strand ssRNA viruses were obtained from NCBI Protein Database. Multiple sequence alignment (MSA) and phylogenetic analysis were performed using MEGA X and PhyML(10). MUSCLE algorithm was used to perform MSA between sequences. Maximum likelihood (ML) model selection was performed using “Smart Model Selection in PhyML” (SMS) (11). Phylogenetic tree construction was performed in PhyML and MEGAX. Generated trees were tested by 100X bootstrapping.

## Results

### Comparing SARS-CoV2 proteome with other RNA viruses reveals significant homologies

SARS-CoV2 is a member of betacoronaviruses, which belong to positive sense ssRNA family of viruses (12). To investigate potential molecular similarities between this virus and other ssRNA viruses, we first compared all of SARS-CoV2 proteins (38 RefSeq proteins, Supp Table 1) with protein sequences from major positive strand ssRNA virus families including Picornaviridae, Flaviviridae, Togaviridae and Caliciviridae. As described in the Methods section, BLASTP and Delta-BLAST algorithms were used to perform the comparisons and best E value/homology scores were determined for each SARS-CoV2 protein. As shown in Figure 1, homology searches showed significant similarities (E values below 1E-30) between SARS-CoV2 non-structural protein 3 (nsp3) as well as orf1a polyprotein (which contains nsp3) with Togavirus proteins (Figure 1a). Far less significant homologies were observed for SARS-CoV2 helicase (E values around 1E-5) with proteins from members of Togaviridae (Figure 1a). When compared with Picornaviridae and Caliciviridae, strongest homologies belonged to SARS-CoV2 helicase, with E values around 1E-3 (Figure 1b, 1c). No significant homologies could be detected with members of Flaviviridae (data not shown). We next compared SARS-CoV2 proteins with major negative strand ssRNA virus families, i.e. Paramyxoviridae, Orthomyxoviridae, Rhabdoviridae and Filoviridae. Homology searches against Paramyxoviridae showed a striking homology between SARS-CoV2 spike protein and fusion proteins of various paramyxoviruses (Figure 1d). Comparison with Orthomyxoviridae and Rhabdoviridae showed very limited homologies (Figure 1e, f), and comparisons with Filoviridae did not show any significant similarities (data not shown). As shown in Figure 2a, observed homologies for nsp3-Togavirus and spike-Paramyxovirus homologies (hereafter named Spike-Fusion) were substantially stronger than other homologies. We then performed homology searches between SARS-CoV2 nsp3 against members of Togaviridae which are known human pathogens. As shown in Figure 2b, Venezuelan equine encephalitis virus (VEEV) followed by rubella virus, Western equine encephalitis virus (WEEV) and Eastern equine encephalitis viruses (EEEV) showed highest log E value-scores. Similar comparisons were done for SARS-CoV2 spike versus Paramyxoviridae which are human pathogens. Here, measles virus followed by henipaviruses showed greatest homologies (Figure 2c). Delta-BLAST alignments for SARS-CoV2 spike versus measles morbillivirus fusion protein and SARS-CoV2 nsp3 versus VEEV nsp3 are shown in Supplemental Figures 1 and 2. Entire taxonomy report for Delta-BLAST results for nsp3-togavirus and spike-paramyxovirus (i.e. Spike-Fusion) comparisons are shown in Supplemental Tables 3 and 4.

**Figure 1-.**
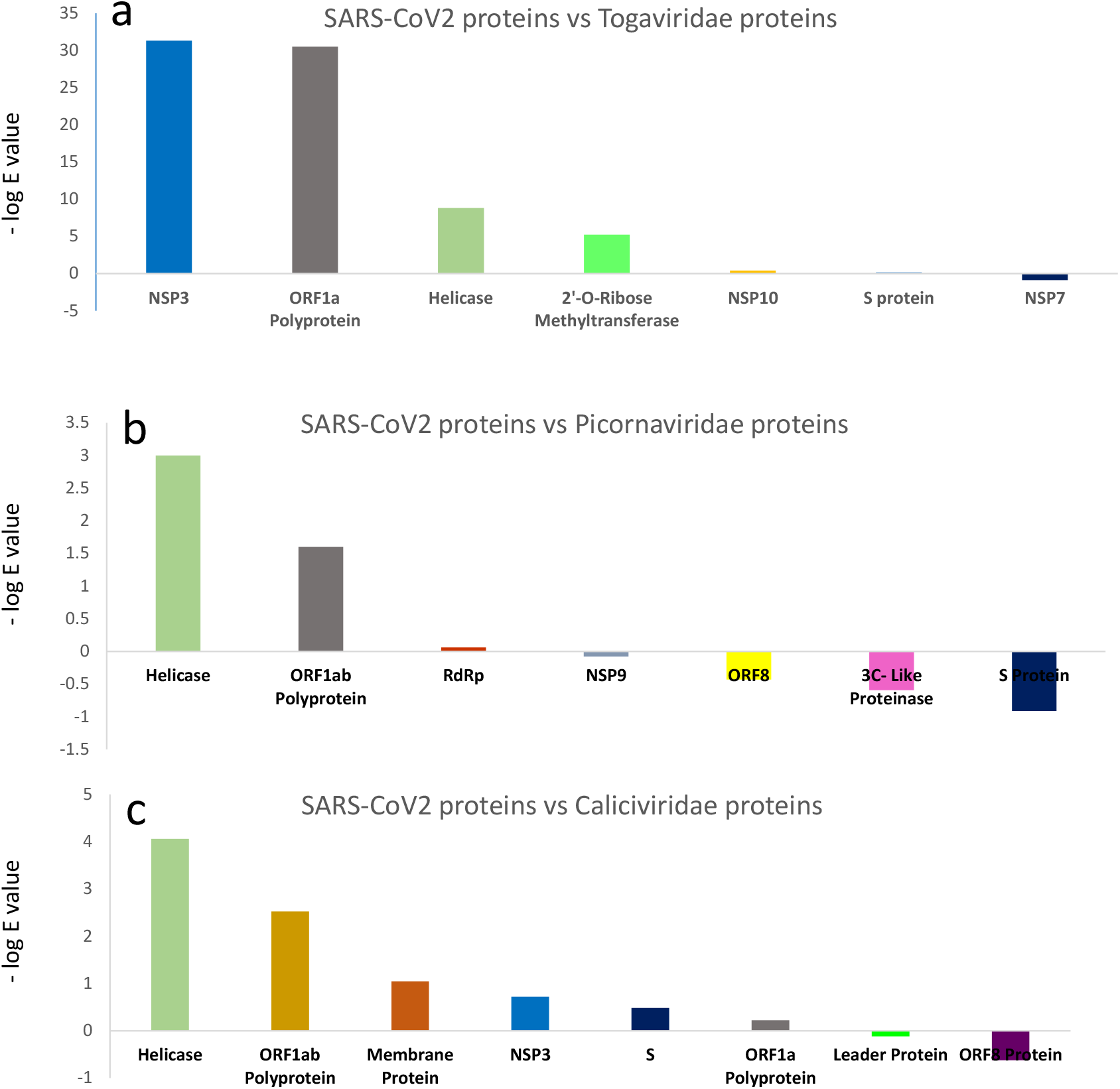
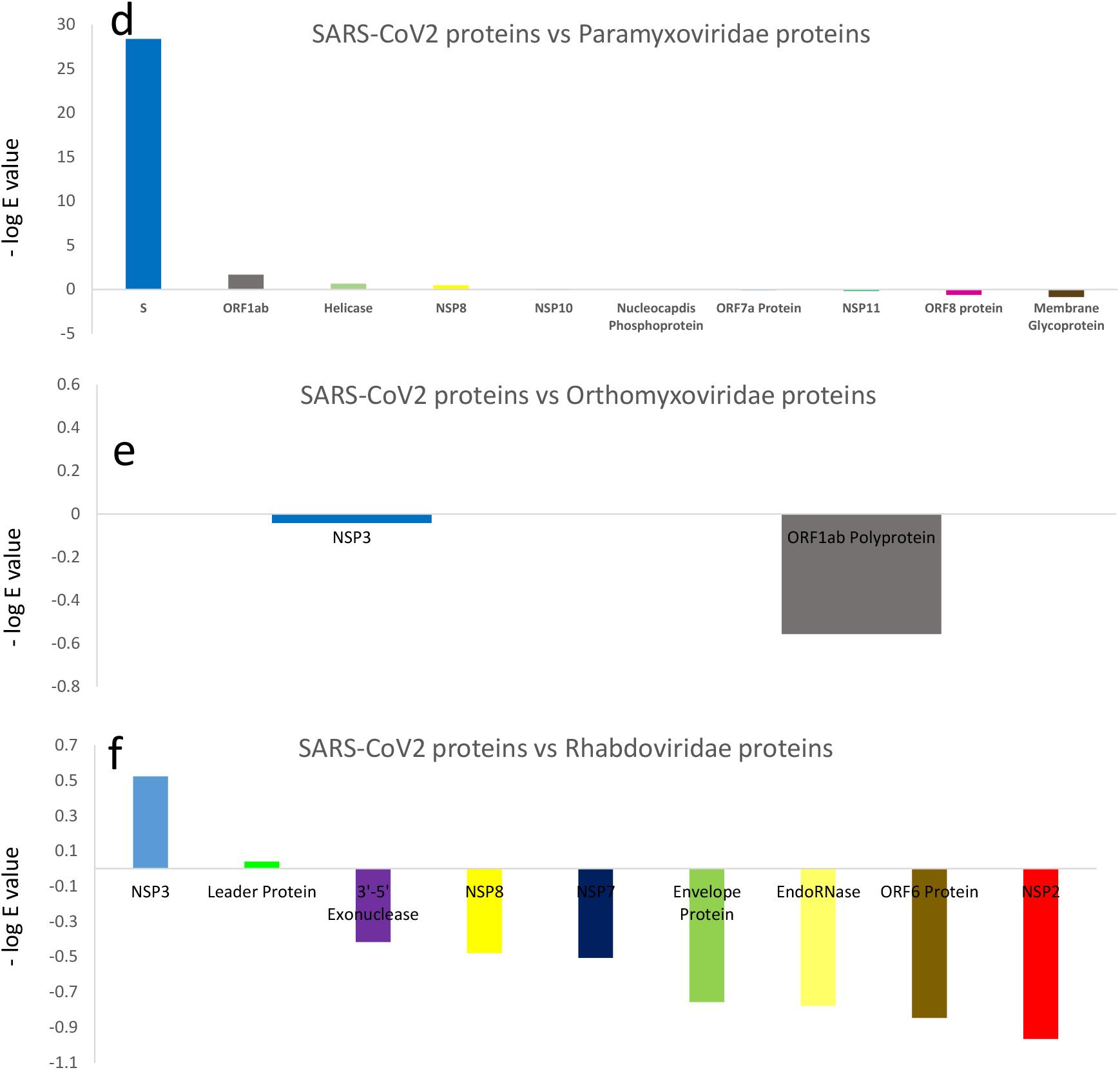
Homology between SARS-CoV2 proteins and major ssRNA virus families. Results of homology searches for 38 SARS-CoV2 RefSeq proteins against major positive strand and negative strand ssRNA viruses are shown as bar graphs. Y axis in each graph shows highest negative log E value obtained for each protein query.

**Figure 2-.**
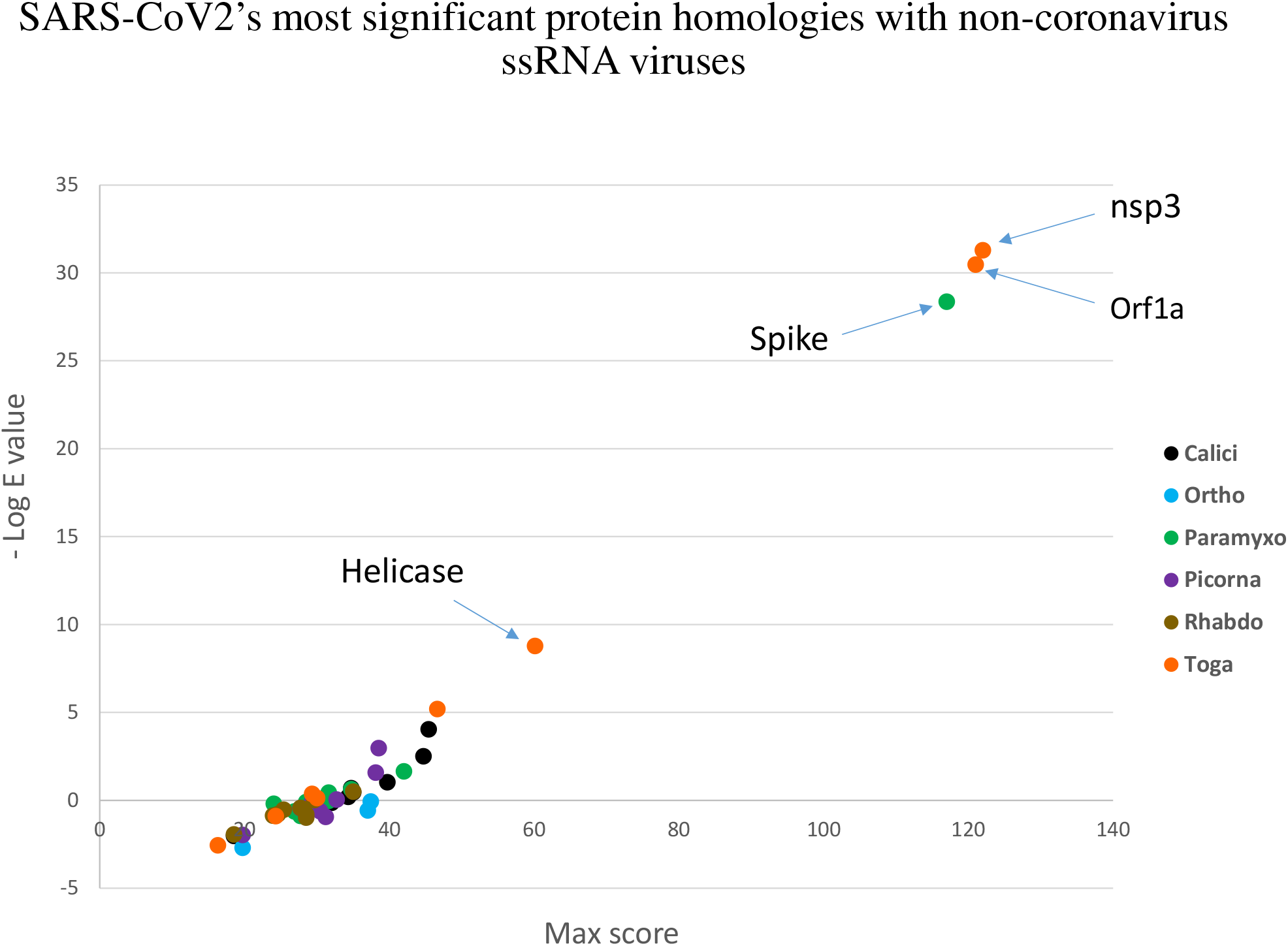
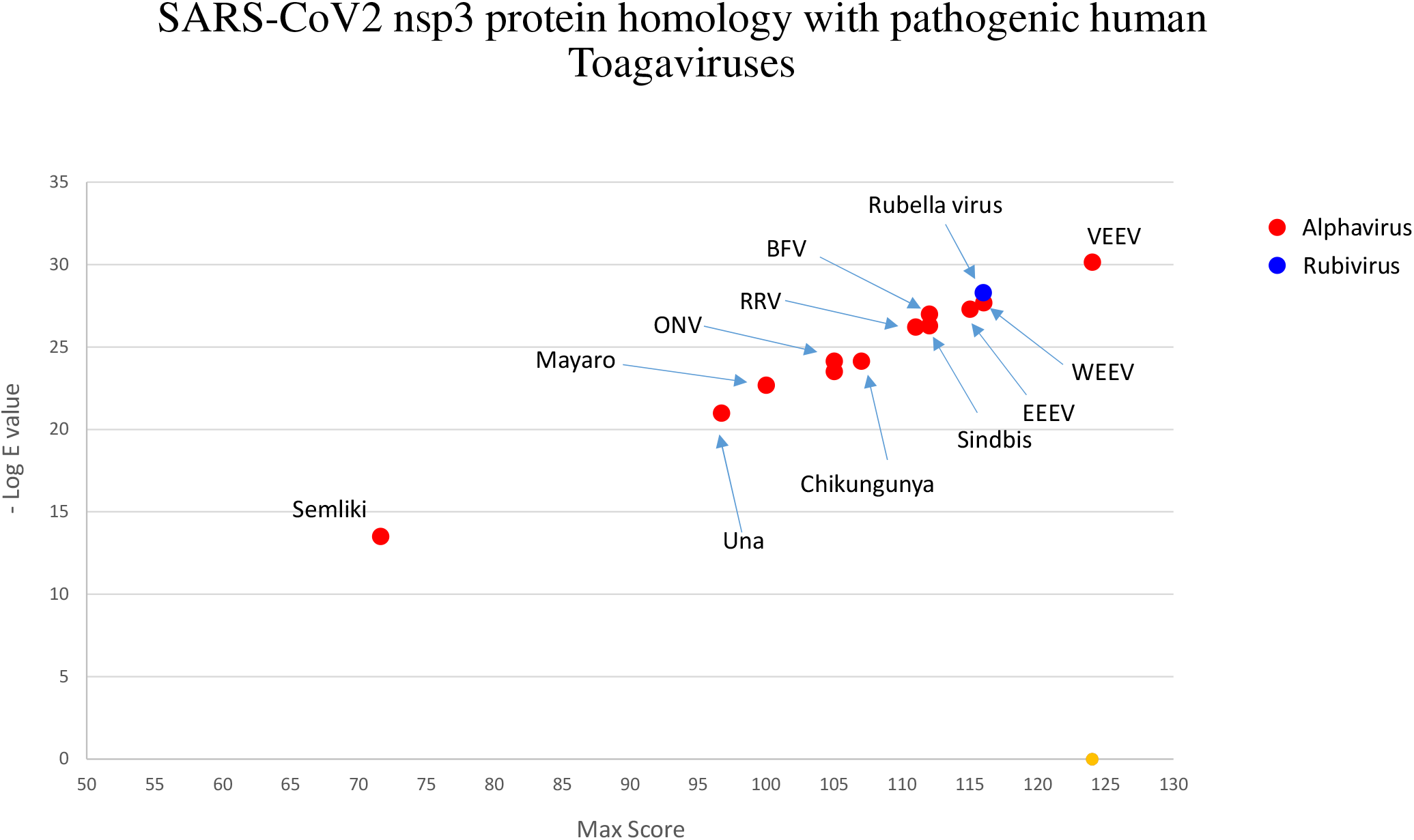
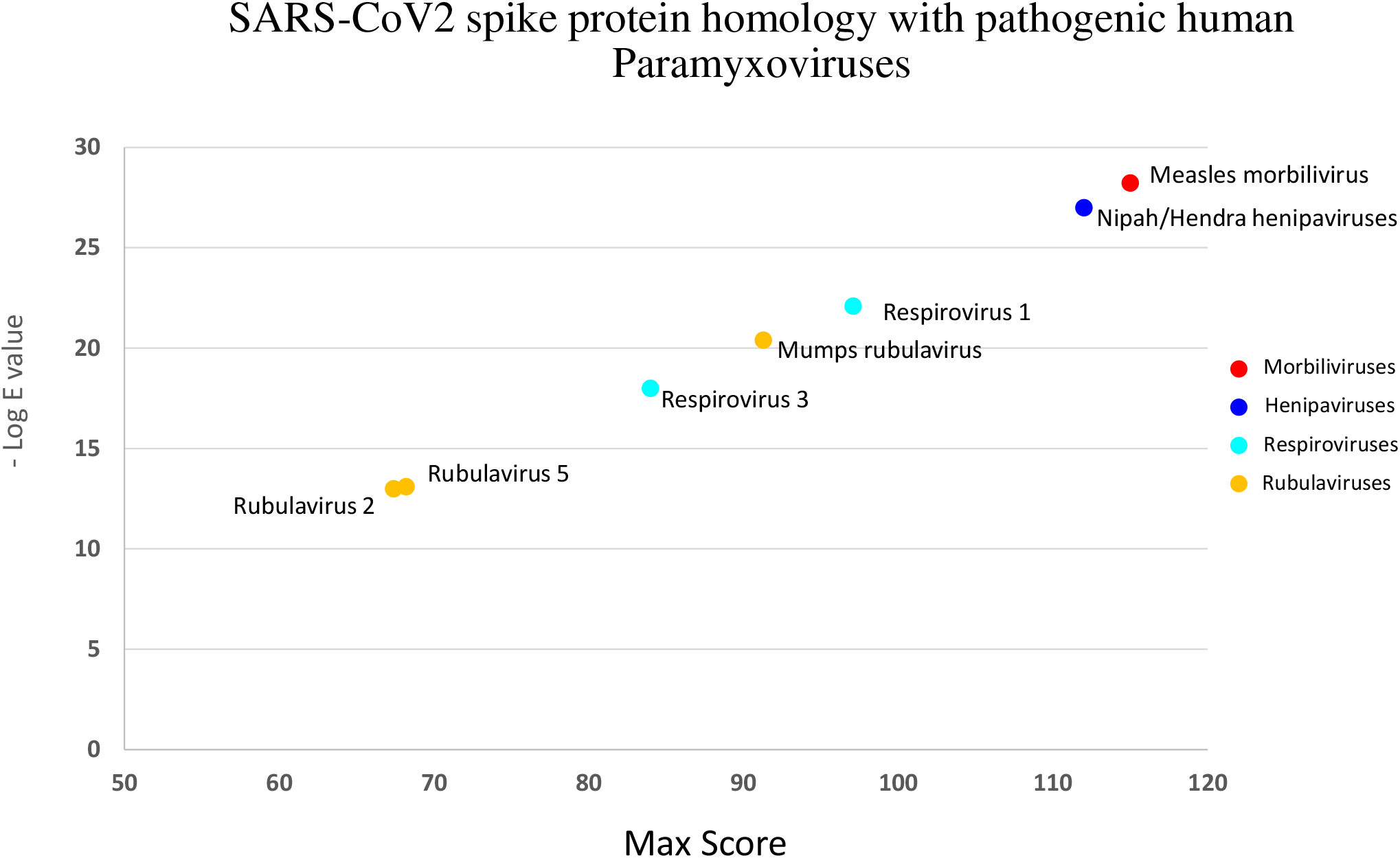
Protein homologies for all SARS-CoV2 proteins against 6 groups of ssRNA viruses. Score-negative log E value for various SARS-CoV2 proteins against ssRNA virus families. For each protein query, only the highest Score-negative log E value is shown (a). Individual score-negative log E values for SARS-CoV2 nsp3 protein versus human pathogenic members of Togaviridae (b). Individual score-negative log E values for SARS-CoV2 spike protein versus human pathogenic members of Paramyxoviridae (c).

### Coronaviridae spike proteins show different degrees of homology with paramyxoviruses

Considering that coronavirus spike proteins are important players in cell entry as well as anti-viral immune responses, we focused the rest of this work on spike proteins. We asked whether the observed similarity between spike protein and paramyxovirus fusion proteins is limited to SARS-CoV2 or that it is a general phenomenon among coronaviruses. To answer this question, we compiled a list of viruses representative of alpha, beta, gamma and delta coronaviruses and obtained their spike protein sequences from NCBI protein database (Supplemental Table 2). We next performed Delta-BLAST searches for each of these spike proteins against Paramyxoviridae protein sequences. Our analyses showed that Spike-Fusion homologies do exist for some other beta and alpha coronaviruses and to a much more limited degree for gamma and delta coronaviruses (Figure 3). Before the emergence of SARS-CoV2, pathogenic human betacoronaviruses included highly pathogenic SARS-CoV and MERS-CoV as well as far less pathogenic OC43 and HKU1. When compared with paramyxoviruses, two highly pathogenic coronaviruses (i.e. SARS-CoV2 and SARS-CoV) showed a much higher degree of homology with paramyxovirus surface proteins, compared with homologies observed for MERS-CoV, HKU1 and OC43 (Figure 3, red bars). Of note, SARS-CoV2’s spike homology was much stronger (E value ~ 1E-28) than SARS-CoV (E value ~ 1E-9). One animal coronavirus, i.e. Bat CoV BM48 also showed a high degree of similarity with Paramyxoviridae fusion proteins, albeit with a less significant E value compared with SARS-CoV2 (E value ~ 1E-15).

**Figure 3.**
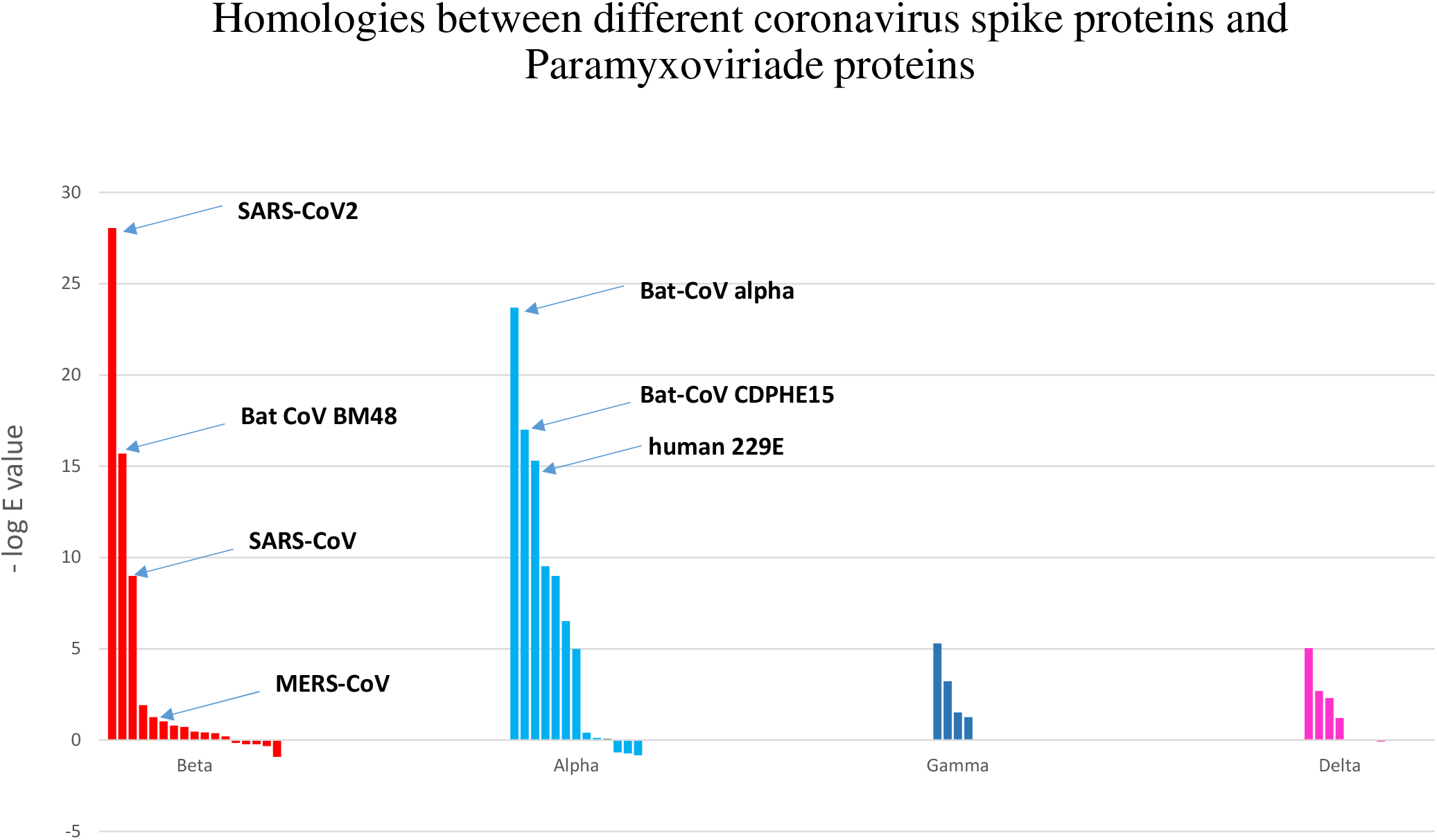
Spike proteins from different coronaviruses show different degrees of homology with paramyxovirus fusion proteins. Negative log E values are shown for spike-paramyxovirus comparisons for members of alpha, beta, gamma and delta coronaviruses. The complete list of these viruses as well as their spike protein accession numbers are shown in Supplemental Table 2.

Among alphacoronaviruses, strongest homologies belonged to alpha-Bat CoV followed by CDPHE15 Bat CoV and human 229E (Figure 3, blue bars). Coronaviruses express three proteins at their surface; S (spike), E (envelope) and M (membrane). To determine whether homologies might also exist for the latter two proteins, we performed similar analyses for all coronavirus E and M proteins against paramyxoviruses. No significant similarities with paramyxoviruses were observed for either E or M proteins (data not shown).

### Spike-paramyxovirus homology regions might bear immunological significance

Protein sequence similarities between different pathogens might have phylogenetic, pathogenic and immunological implications. Considering that spike protein is a target for humoral immune responses, we asked whether observed homology between SARS-CoV2 spike and paramyxoviruses might have any potential immunological importance. Considering that it is a recently emerged pathogen, SARS-CoV2 protein epitopes are not yet fully known. However, a substantial body of experimental work has been performed on related SARS-CoV antigens. To find immunologically important regions in SARS-CoV2 spike protein, we searched Immune Epitope Database (IEDB) using SARS-CoV2 spike protein sequence. We looked for exact substring matches in spike protein with experimentally verified B cell, T cell or MHC binding epitopes (Supplemental Tables 5-7). As shown in Table 1, 33 B cell, 9 T cell and 65 MHC binding exact epitope matches were found in SARS-CoV2 spike protein. We next determined the distribution of these epitopes inside spike protein, using pairwise BLASTP search. As shown in Figure 4a, most B cell epitopes were concentrated in two regions of spike protein; a stretch of amino acids from 950-1050 and another stretch from 1150 to 1200. This first stretch coincided with ‘Spike-Fusion’ homology region. Similarly, when we looked at the distribution of T cell epitopes, 5 out of 9 IEDB T cell epitopes mapped within the homology region (Figure 4b). Of MHC binding peptides, 23 out of 65 peptide were located inside homology region. Overall, these findings point to potential importance of Spike-Fusion homology region in stimulating humoral and cellular immune responses.

**Table 1-.** Distribution of B cell, T cell and MHC binding epitopes in SARS-CoV2 spike protein. Table shows the total number of B cell, T cell and MHC binding epitopes that corresponded with substrings of SARS-CoV2 spike protein from IEDB, as well as the number of epitopes present in the Spike-Fusion homology region.

| Type of epitope<br>(experimentally<br>verified IEDB<br>epitopes) | Total number from<br>IEDB | Number of Aligned<br>epitopes (E value <0.1) | Number of epitopes<br>inside Spike-Fusion<br>homology region |
| --- | --- | --- | --- |
| B cell epitopes | 33 | 29 | 10 |
| T cell epitopes | 9 | 9 | 5 |
| MHC binding<br>epitopes | 65 | 64 | 23 |

**Figure 4.**
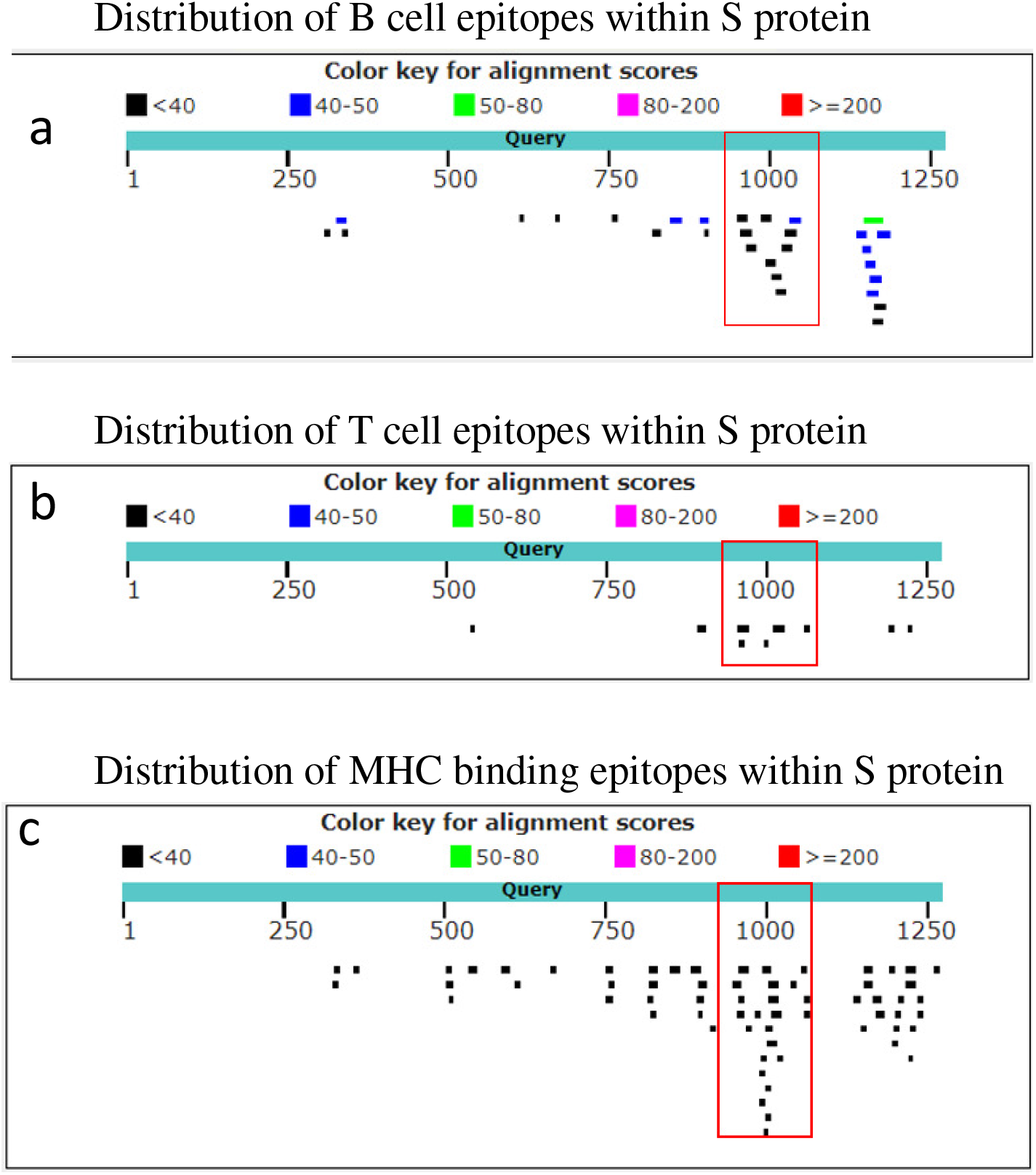
Distribution of IEDB B cell, T cell and MHC binding epitopes inside SARS-CoV2 spike protein. Experimentally verified epitopes which matched substrings of SARS-CoV2 spike protein were obtained from IEDB and aligned with spike protein sequence to determine their distribution across the protein.

### Observed SARS-CoV2 Spike-Fusion homologies do not follow the general pattern of ssRNA virus phylogenetic relations

The most likely explanation for similarities between nucleic acid or protein sequences from various microorganisms is ‘shared ancestry’. To examine whether observed similarities between SARS-CoV2 spike and paramyxovirus surface proteins are a function of shared ancestry, we performed phylogenetic analyses on various members of ssRNA viruses. The protein sequence of RNA-dependent RNA polymerase (RdRP) is widely used for performing phylogenetic studies on RNA viruses. Representative pathogenic viruses were selected from each group (shown in Supplemental Table 8) and relevant RdRP sequences were obtained from NCBI Protein Database. Phylogenetic analysis using Maximum Likelihood method showed separate clusters for the majority of positive strand ssRNA versus negative strand ssRNA viruses (Figure 5). Of note, members of Coronaviridae were clustered separately from Paramyxoviridae based on RdRP protein sequences. This indicated that observed similarity between spike-surface proteins were not a reflection of general phylogenetic similarity. Phylogenetic analyses using distance-based methods also led to similar results (data not shown).

**Figure 5.**
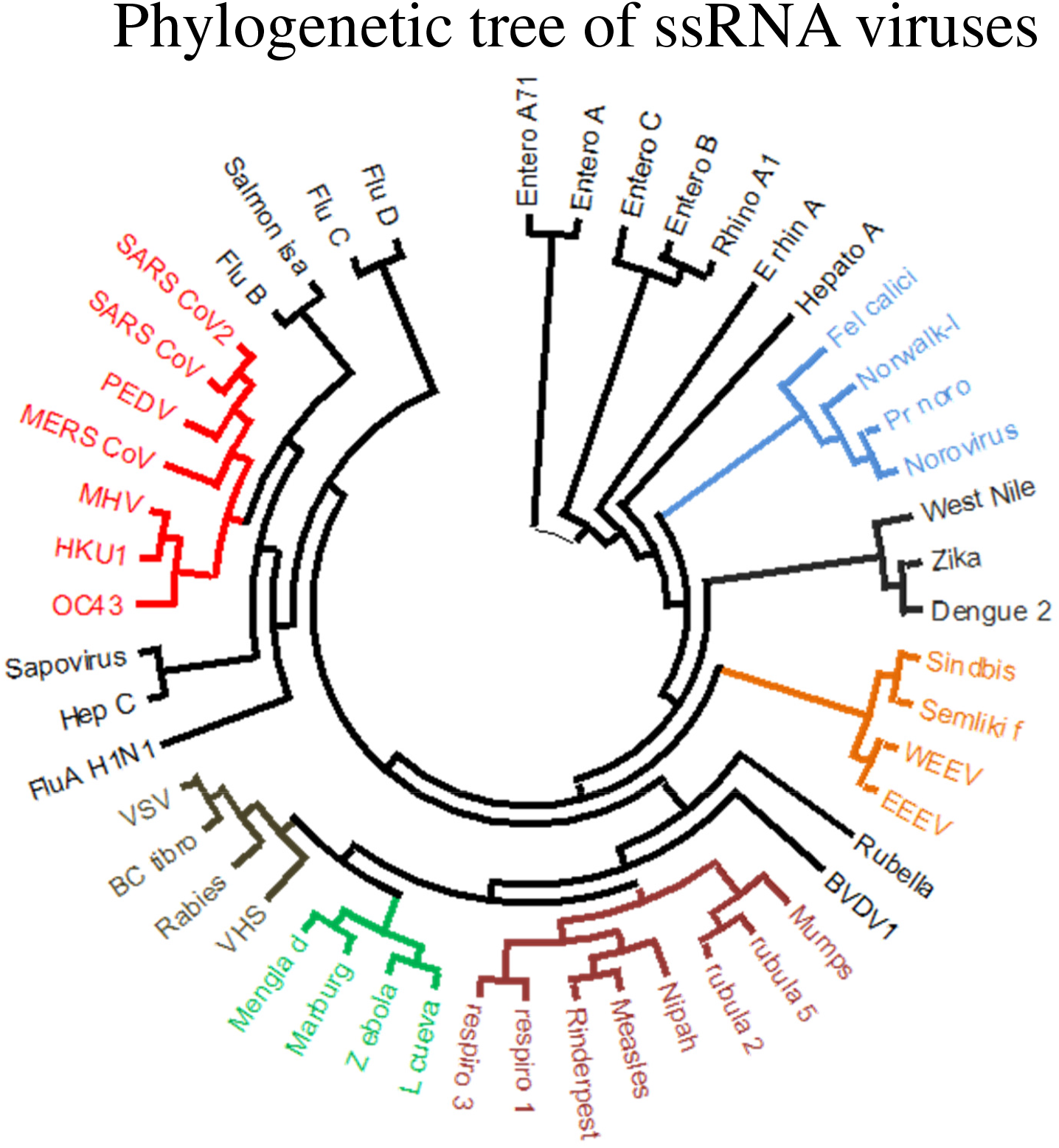
Phylogenetic tree of ssRNA viruses. Phylogenetic tree of representative members of positive sense strand ssRNA viruses (i.e. Picornaviridae, Caliciviridae, Coronaviridae, Flaviviridae and Togaviridae) and negative sense strand ssRNA viruses (i.e. Orthomyxoviridae, Paramyxoviridae, Filoviridae, Rhabdoviridae) was drawn based on viral RdRP protein sequences. Virus name abbreviations: BC tibro, Bas-Congo tibrovirus; BVDV1, Bovine viral diarrhea virus 1; EEEV, Eastern equine encephalitis virus; L Cueva, Lloviu cuevavirus; PEDV, Porcine epidemic diarrhea virus; Pr-nor, Primate norovirus; Semliki F, Semliki Forest virus; VHS, Viral hemorrhagic septicemia virus; VSV, Vesicular stomatitis Indiana virus; WEEV, Western equine encephalitis virus.

## Discussion

Microbial pathogens display different degrees of likeness at the level of their macromolecules, as well their biological behavior. This is generally a consequence of shared ancestry, horizontal gene transfer or convergent evolution. When it comes to infectious diseases, these similarities might bear epidemiological or clinical importance. This is especially so when a new pathogen appears, whose evolution, pathogenesis and immunology is largely unknown. In this bioinformatic study, we found a potentially important homology between SARS-CoV2 spike protein and paramyxoviral surface proteins. This homology was far more significant than any other homologies between SARS-CoV2 and ssRNA viruses (except for SARS-CoV2 nsp3 homology with Togaviruses). Although this homology also existed for some other pathogenic betacoronavirus spike proteins, the one observed for SARS-CoV2 demonstrated a much higher score-log E value. When analyzed for the presence of experimentally verified epitopes (which are mostly derived from studies on SARS-CoV proteins), a high number of B cell, T cell and MHC binding epitopes were observed inside this Spike-Fusion homology region. Hence, it seemed that this observed homology was likely more than a coincidental and biologically inert similarity.

Paramyxoviruses include several major human and animal pathogens, with well-recognized pathophysiology and immune behavior (13–16). Measles and mumps viruses are known human paramyxoviruses and canine distemper and Rinderpest viruses are animal pathogens. In our analyses, a reptilian paramyxovirus followed by canine, rinderpest and measles morbiliviruses revealed the highest degrees of homology with SARS-CoV2 spike protein. Among human pathogens, lower levels of homology were seen with fusion proteins from henipaviruses and respirovirus 1. The observed Spike-Fusion protein homology was not limited to SARS-CoV2. Spike proteins from other betacoronaviruses (e.g. bat CoV BM48 and SARS-CoV) as well as some alpha coronaviruses also showed some levels of homology. However, when compared in terms of expect value, SARS-CoV2 spike-fusion homology was orders of magnitude stronger that other pathogenic human coronaviruses, i.e. SARS-CoV and MERS-CoV.

Evolution from a common ancestor is usually the underlying reason for the existence of homologies between nucleic acid and/or protein sequences. Indeed, the concept of “sequence homology” entails “shared ancestry” and sequence similarities between taxa which do not share a recent common ancestor are referred to as ‘homoplasy’ (17, 18). Phylogenetic relations of RNA viruses is usually inferred based on analyses of well conserved proteins including RNA polymerases. RdRP-based phylogenetic analysis of representative members of ssRNA viruses showed that the observed Spike-Fusion protein similarity does not seem to follow the branching pattern of inferred phylogenetic tree. While various RNA viruses ought to share a common ancestor at some level of evolution, our current analysis suggests that factors other than shared ancestry might be at work. Presumably, there might be a degree of convergent evolution between SARS-CoV2 spike and paramyxovirus fusion proteins, which might be driven by selection pressure from host-pathogen interactions. Regardless of the underlying evolutionary mechanism, similar viral protein sequences might bear immunological importance, especially if they are observed in proteins which can be readily targeted by adaptive immune responses. Studies on previous coronaviruses have shown that spike protein is a target for humoral as well as cellular immune responses (19–21). When analyzed for the presence of experimentally verified B cell and/or cell epitopes Spike-Fusion homology region showed a high concentration of these epitopes, compared with the rest the protein sequence. As alluded to in the introduction, some recent reports have suggested that vaccination using live MMR vaccine might offer a degree of protection against COVID-19 infection. Consistent with our data, a pre-print study by Franklin et al has shown protein sequence as well as 3D structure similarities between SARS-CoV2 spike and measles/mumps virus fusion proteins (7). They have also reported epidemiological data indicating that MMR vaccination is indeed associated with lower COVID-19 frequency in different countries (7). It should be noted that, while antigenic similarities raise the possibility of cross-protective adaptive immune responses, this does not exclude the possibility of involvement of innate immune mechanisms, as shared pathogenic molecular structures might also lead to similar patterns of innate immune activation.

Unraveling different aspects of the evolution of microbial pathogens is of paramount importance in predicting, understanding and controlling infectious disease epidemics. Ordinary bioinformatic analyses on nucleic acid and/or proteins might give us some hints about the relatedness or dissimilarities between pathogenic microorganisms, but grasping the complexity of this dynamism would likely require integrated analyses of host(s)-pathogen(s) co-evolution.

