## Supplemental Figure 1 for "SARS-CoV2 spike protein displays biologically significant similarities with paramyxovirus surface proteins; a bioinformatics study"

Supp Figure 1- Delta-BLAST results for SARS-CoV2 spike protein (Query) versus measles mobilivirus fusion protein (Subject)

### fusion protein [Measles morbillivirus]

**Sequence ID:** NP_056922.1 **Length:** 550 **Number of Matches:** 1

**Range 1: 119 to 249** GenPept Graphics ▼ Next Match ▲ Previous Match

| Score | Expect | Method | Identities | Positives | Gaps |
|---|---|---|---|---|---|
| 115 bits(289) | 4e-28 | Composition-based stats. | 33/139(24%) | 55/139(39%) | 24/139(17%) |

```
Query  890   AGAALQIPFAMQMAYRFNGIGVTQNVLYENQKLIANQFNS------AIGKIQDSLSSTAS  943
             AGAAL +  A Q+     GI + Q++L  N + I N   S      AI  I+ +
Sbjct  119   AGAALGVATAAQIT---AGIALHQSML--NSQAIDNLRASLETTNQAIEAIRQAGQEMIL  173

Query  944   ALGKLQDVVNQN-AQALNTLVKQLSSNFGAISSVLNDILSRL---------DKVEAEVQI  993
             A+  +QD +N     ++N L   L      +   L    + +         D + AE+ I
Sbjct  174   AVQGVQDYINNELIPSMNQLSCDLIGQKLGLK--LLRYYTEILSLFGPSLRDPISAEISI  231

Query  994   DRLITGRLQSLQTYVTQQL  1012
                ++   L      V ++L
Sbjct  232   Q-ALSYALGGDINKVLEKL  249
```
