## Supplemental Figure 2 for "SARS-CoV2 spike protein displays biologically significant similarities with paramyxovirus surface proteins; a bioinformatics study"

### Supp Figure 2- Delta-BLAST results for SARS-CoV2 nsp3 protein (Query) versus Venezuelan equine encephalitis virus nsp3 protein (Subject)

#### putative nonstructural protein nsP3 [Venezuelan equine encephalitis virus]

**Sequence ID:** NP_740698.1 **Length:** 557 **Number of Matches:** 1

**Range 1: 6 to 165** GenPept Graphics Next Match Previous Match

| Score | Expect | Method | Identities | Positives | Gaps |
| --- | --- | --- | --- | --- | --- |
| 124 bits(310) | 1e-30 | Composition-based stats. | 49/179(27%) | 79/179(44%) | 34/179(18%) |

```
Query  222  IKNADIVEEAKKVKPTVVVNAANVYLKHGGGVAGALNKATNNAMQVESDDYIATNGPLKV  281
            +   DI          V++NAAN   + GGGV GAL K    +  +          P++V
Sbjct  6    VVRGDIAT----ATEGVIINAANSKGQPGGGVCGALYKKFPESFDL---------QPIEV  52

Query  282  GGSCVLSGHNLAKHCLHVVGPNVNKGEDIQ---LLKSAYENFNQ------HEVLLAPLLS  332
            G + ++ G   AKH +H VGPN NK  +++    L  AYE+  +      ++ +  PLLS
Sbjct  53   GKARLVKGA--AKHIIHAVGPNFNKVSEVEGDKQLAEAYESIAKIVNDNNYKSVAIPLLS  110

Query  333  AGIFGADPI---HSLR---VCVDTVRTNVYLAVFDKNLYDKLVSSFLEMKSEKQVEQKI  385
             GIF  +      SL      +DT   +V +   D     K   +  E  + ++  ++I
Sbjct  111  TGIFSGNKDRLTQSLNHLLTALDTTDADVAIYCRD----KKWEMTLKEAVARREAVEEI  165
```
