## Supplemental Table 1 for "SARS-CoV2 spike protein displays biologically significant similarities with paramyxovirus surface proteins; a bioinformatics study"

### Supp Table 1- SARS-CoV2 RefSeq protein accession numbers and names

| No | List of SARS-CoV2 Accession Numbers | Name | No | List of SARS-CoV2 Accession Numbers | Name |
|---|---|---|---|---|---|
| 1 | YP_009742617.1 | nsp10 | 20 | YP_009725304.1 | nsp8 |
| 2 | YP_009742616.1 | nsp9 | 21 | YP_009725303.1 | nsp7 |
| 3 | YP_009742615.1 | nsp8 | 22 | YP_009725302.1 | nsp6 |
| 4 | YP_009742614.1 | nsp7 | 23 | YP_009725301.1 | 3C-like proteinase |
| 5 | YP_009742613.1 | nsp6 | 24 | YP_009725300.1 | nsp4 |
| 6 | YP_009742612.1 | 3C-like proteinase | 25 | YP_009725299.1 | nsp3 |
| 7 | YP_009742611.1 | nsp4 | 26 | YP_009725298.1 | nsp2 |
| 8 | YP_009742610.1 | nsp3 | 27 | YP_009725297.1 | leader protein |
| 9 | YP_009742609.1 | nsp2 | 28 | YP_009725295.1 | orf1a polyprotein |
| 10 | YP_009742608.1 | leader protein | 29 | YP_009725255.1 | ORF10 protein |
| 11 | YP_009725318.1 | ORF7b | 30 | YP_009724397.2 | nucleocapsid phosphoprotein |
| 12 | YP_009725312.1 | nsp11 | 31 | YP_009724396.1 | ORF8 protein |
| 13 | YP_009725311.1 | 2'-O-ribose methyltransferase | 32 | YP_009724395.1 | ORF7a protein |
| 14 | YP_009725310.1 | endoRNAse | 33 | YP_009724394.1 | ORF6 protein |
| 15 | YP_009725309.1 | 3'-to-5' exonuclease | 34 | YP_009724393.1 | membrane glycoprotein |
| 16 | YP_009725308.1 | helicase | 35 | YP_009724392.1 | envelope protein |
| 17 | YP_009725307.1 | RNA-dependent RNA polymerase | 36 | YP_009724391.1 | ORF3a protein |
| 18 | YP_009725306.1 | nsp10 | 37 | YP_009724390.1 | surface glycoprotein |
| 19 | YP_009725305.1 | nsp9 | 38 | YP_009724389.1 | orf1ab polyprotein |
