## Supplemental Table 2 for "SARS-CoV2 spike protein displays biologically significant similarities with paramyxovirus surface proteins; a bioinformatics study"

### Supp Table 2- Spike protein accession numbers for various coronaviruses

| Genera | Host | Organism | SYM | Accession number |
|---|---|---|---|---|
| Alpha | Human | Human coronavirus 229E | HC229 | NP_073551.1 |
| Alpha | Human | Human coronavirus NL63 | HCN | YP_003767.1 |
| Alpha | Animal | Alphacoronavirus Bat-CoV | ABC | YP_009755890.1 |
| Alpha | Animal | Camel alphacoronavirus | CA | YP_009194639.1 |
| Alpha | Animal | Feline infectious peritonitis virus | FIPV | YP_004070194.1 |
| Alpha | Animal | Shrew coronavirus | SC | YP_009755839.1 |
| Alpha | Animal | Wencheng Sm shrew coronavirus | WSC | YP_009389425.1 |
| Alpha | Animal | Coronavirus AcCoV-JC34 | CAJ | YP_009380521.1 |
| Alpha | Animal | Swine enteric coronavirus | SEC | YP_009199242.1 |
| Alpha | Animal | Mink coronavirus strain WD1127 | MCS | YP_009019182.1 |
| Alpha | Animal | Porcine epidemic diarrhea virus | PEDV | NP_598310.1 |
| Alpha | Animal | Scotophilus bat coronavirus 512 | SBC | YP_001351684.1 |
| Alpha | Animal | Bat coronavirus CDPHE15 | BCC | YP_008439202.1 |
| Beta | Human | Severe acute respiratory syndrome coronavirus 2 | SARS-2 | YP_009724390.1 |
| Beta | Human | Middle East respiratory syndrome-related coronavirus | MERS | YP_009047204.1 |
| Beta | Human | Human coronavirus HKU1 | HCH | YP_173238.1 |
| Beta | Human | Severe acute respiratory syndrome-related coronavirus | SARS | NP_828851.1 |
| Beta | Human | Human coronavirus OC43 | HCO | YP_009555241.1 |
| Beta | Animal | Betacoronavirus Erinaceus | BE | YP_009513010.1 |
| Beta | Animal | Betacoronavirus England 1 | BE-1 | YP_007188579.1 |
| Beta | Animal | Rabbit coronavirus HKU14 | RCH | YP_005454245.1 |
| Beta | Animal | Bat coronavirus BM48 | BCB | YP_003858584.1 |
| Beta | Animal | Rousettus bat coronavirus HKU9 | RBCH | YP_001039971.1 |
| Beta | Animal | Pipistrellus bat coronavirus HKU5 | PBCH | YP_001039962.1 |
| Beta | Animal | Tylonycteris bat coronavirus HKU4 | TBCH | YP_001039953.1 |
| Beta | Animal | Bovine coronavirus | BC | NP_150077.1 |
| Beta | Animal | Murine hepatitis virus | MHV | NP_045300.1 |
| Beta | Animal | Betacoronavirus HKU24 | BH | YP_009113025.1 |
| Beta | Animal | Bat Hp-betacoronavirus/Zhejiang2013 | BHB | YP_009072440.1 |
| Beta | Animal | Rat coronavirus Parker | RCP | YP_003029848.1 |
| Delta | Animal | Porcine coronavirus HKU15 | PCH | YP_009513021.1 |
| Delta | Animal | Sparrow coronavirus HKU17 | SCH | YP_005352846.1 |
| Delta | Animal | White-eye coronavirus HKU16 | WECH | YP_005352838.1 |
| Delta | Animal | Common moorhen coronavirus HKU21 | CMCH | YP_005352881.1 |
| Delta | Animal | Wigeon coronavirus HKU20 | WCH | YP_005352871.1 |
| Delta | Animal | Night heron coronavirus HKU19 | NHCH | YP_005352863.1 |
| Delta | Animal | Magpie-robin coronavirus HKU18 | MRCH | YP_005352854.1 |
| Delta | Animal | Munia coronavirus HKU13-3514 | MCH | YP_002308506.1 |
| Gamma | Animal | Turkey coronavirus | TC | YP_001941166.1 |
| Gamma | Animal | Infectious bronchitis virus | IBV | NP_040831.1 |
| Gamma | Animal | Canada goose coronavirus | CGC | YP_009755897.1 |
| Gamma | Animal | Beluga whale coronavirus SW1 | BWC | YP_001876437.1 |
