## Supplemental Table 3 for "SARS-CoV2 spike protein displays biologically significant similarities with paramyxovirus surface proteins; a bioinformatics study"

### Supp Table 3- Taxonomy report for members of Toagaviridae which show homology with SARS-CoV2 nsp3 protein (first 500 hits, E values below 1E-24)

| Taxonomy Report | | | |
|---|---|---|---|
| Taxonomy | Number of hits | Number of Organisms | Description |
| Alphavirus | 678 | 26 | |
| . Venezuelan equine encephalitis virus | 154 | 6 | Venezuelan equine encephalitis virus hits |
| .. Venezuelan equine encephalitis virus (strain P676) | 9 | 1 | Venezuelan equine encephalitis virus (strain P676) hits |
| .. Venezuelan equine encephalitis virus (strain Mena II) | 1 | 1 | Venezuelan equine encephalitis virus (strain Mena II) hits |
| .. Venezuelan equine encephalitis virus (strain 3880) | 1 | 1 | Venezuelan equine encephalitis virus (strain 3880) hits |
| .. Venezuelan equine encephalitis virus (strain Trinidad donkey) | 1 | 1 | Venezuelan equine encephalitis virus (strain Trinidad donkey) hits |
| .. Venezuelan equine encephalitis virus (strain CPA201) | 1 | 1 | Venezuelan equine encephalitis virus (strain CPA201) hits |
| . Fort Morgan virus | 3 | 2 | Fort Morgan virus hits |
| .. Buggy Creek virus | 1 | 1 | Buggy Creek virus hits |
| . Tonate virus | 3 | 1 | Tonate virus hits |
| . Mucambo virus | 4 | 1 | Mucambo virus hits |
| . Cabassou virus | 3 | 1 | Cabassou virus hits |
| . Rio Negro virus | 3 | 1 | Rio Negro virus hits |
| . Mosso das Pedras virus | 3 | 1 | Mosso das Pedras virus hits |
| . Everglades virus | 3 | 1 | Everglades virus hits |
| . Pixuna virus | 3 | 1 | Pixuna virus hits |
| . Whataroa virus | 3 | 1 | Whataroa virus hits |
| . Western equine encephalitis virus | 46 | 1 | Western equine encephalitis virus hits |
| . Highlands J virus | 13 | 1 | Highlands J virus hits |
| . Chikungunya virus | 4 | 2 | Chikungunya virus hits |
| .. Chikungunya virus strain S27-African prototype | 12 | 1 | Chikungunya virus strain S27-African prototype hits |
| . Eastern equine encephalitis virus | 304 | 1 | Eastern equine encephalitis virus hits |
| . Ross River virus | 97 | 1 | Ross River virus hits |
| . Barmah Forest virus | 1 | 1 | Barmah Forest virus hits |
| . Sindbis virus | 1 | 1 | Sindbis virus hits |
| . Aura virus | 1 | 1 | Aura virus hits |
| . Bebaru virus | 3 | 1 | Bebaru virus hits |
