## Supplemental Table 4 for "SARS-CoV2 spike protein displays biologically significant similarities with paramyxovirus surface proteins; a bioinformatics study"

### Supp Table 4- Taxonomy report for members of Paramyxoviridae which show homology with SARS-CoV2 spike protein (first 500 hits, E values below 1E-24)

Taxonomy Report

| Taxonomy | Number of hits | Number of Organisms | Description |
|---|---|---|---|
| Paramyxoviridae | 770 | 51 | |
| . Reptilian paramyxovirus | 5 | 1 | Reptilian paramyxovirus hits |
| . Orthoparamyxovirinae | 765 | 50 | |
| .. Morbillivirus | 695 | 43 | |
| ... Canine morbillivirus | 303 | 2 | Canine morbillivirus hits |
| .... Canine distemper virus strain Onderstepoort | 2 | 1 | Canine distemper virus strain Onderstepoort hits |
| ... Rinderpest morbillivirus | 104 | 3 | Rinderpest morbillivirus hits |
| .... Rinderpest virus (strain L) | 1 | 1 | Rinderpest virus (strain L) hits |
| .... Rinderpest virus (strain rbt1) | 1 | 1 | Rinderpest virus (strain rbt1) hits |
| ... Measles morbillivirus | 104 | 37 | Measles morbillivirus hits |
| .... Measles virus genotypes and isolates | 162 | 22 | |
| ..... Measles virus genotype B3 | 15 | 2 | Measles virus genotype B3 hits |
| ...... Measles virus genotype B3.1 | 1 | 1 | Measles virus genotype B3.1 hits |
| ..... Measles virus genotype H1 | 69 | 1 | Measles virus genotype H1 hits |
| ..... Measles virus genotype D8 | 46 | 1 | Measles virus genotype D8 hits |
| ..... Measles virus genotype D9 | 2 | 1 | Measles virus genotype D9 hits |
| ..... Measles virus genotype A | 1 | 1 | Measles virus genotype A hits |
| ..... Measles virus strain MVi/California.USA/8.04 | 1 | 1 | Measles virus strain MVi/California.USA/8.04 hits |
| ..... Measles virus genotype D4 | 13 | 1 | Measles virus genotype D4 hits |
| ..... Measles virus genotype C2 | 1 | 1 | Measles virus genotype C2 hits |
| ..... Measles virus genotype D11 | 1 | 1 | Measles virus genotype D11 hits |
| ..... Measles virus genotype G3 | 1 | 1 | Measles virus genotype G3 hits |
| ..... Measles virus strain MVi/Texas.USA/4.07 | 1 | 1 | Measles virus strain MVi/Texas.USA/4.07 hits |
| ..... Measles virus strain MVi/California.USA/16.03 | 1 | 1 | Measles virus strain MVi/California.USA/16.03 hits |
| ..... Measles virus strain MVi/Virginia.USA/15.09 | 1 | 1 | Measles virus strain MVi/Virginia.USA/15.09 hits |
| ..... Measles virus strain MVi/Washington.USA/18.08/1 | 1 | 1 | Measles virus strain MVi/Washington.USA/18.08/1 hits |
| ..... Measles virus strain MVi/Arizona.USA/11.08/2 | 1 | 1 | Measles virus strain MVi/Arizona.USA/11.08/2 hits |
| ..... Measles virus genotype G2 | 1 | 1 | Measles virus genotype G2 hits |
| ..... Measles virus strain MVi/New York.USA/26.09/3 | 1 | 1 | Measles virus strain MVi/New York.USA/26.09/3 hits |
| ..... Measles virus strain MVi/Florida.USA/19.09 | 1 | 1 | Measles virus strain MVi/Florida.USA/19.09 hits |
| ..... Measles virus genotype D6 | 1 | 1 | Measles virus genotype D6 hits |
| ..... Measles virus strain MVi/Pennsylvania.USA/20.09 | 1 | 1 | Measles virus strain MVi/Pennsylvania.USA/20.09 hits |
| ..... Measles virus strain MVi/New Jersey.USA/45.05 | 1 | 1 | Measles virus strain MVi/New Jersey.USA/45.05 hits |
| .... Measles virus strain Moraten | 1 | 1 | Measles virus strain Moraten hits |
| .... Measles virus strain Schwarz | 1 | 1 | Measles virus strain Schwarz hits |
