## Supplemental Table 5 for "SARS-CoV2 spike protein displays biologically significant similarities with paramyxovirus surface proteins; a bioinformatics study"

### Supp Table 5- IEDB B cell epitopes with exact matches with substrings of SARS-CoV2 spike protein

| No | Epitope ID | Description | Starting Position | Ending Position | Antigen Name | Antigen Accession | Parent Protein |
|---|---|---|---|---|---|---|---|
| 1 | 30987 | KGIYQTSN | 297 | 304 | Spike glycoprotein precursor | P59594.1 | Spike glycoprotein |
| 2 | 70719 | VRFPNITNLCPFGEVFN | 314 | 330 | E2 glycoprotein precursor | NP_828851.1 | Spike glycoprotein |
| 3 | 15972 | FGEVFNAT | 325 | 332 | Spike glycoprotein precursor | P59594.1 | Spike glycoprotein |
| 4 | 462413 | PLQPE | 361 | 365 | receptor tyrosine-protein kinase erbB-2 isoform e | NP_001276867.1 | Receptor tyrosine-protein kinase erbB-2 |
| 5 | 52020 | QQFGRD | 549 | 554 | Spike glycoprotein precursor | P59594.1 | Spike glycoprotein |
| 6 | 558455 | LYQDVN | 597 | 602 | Spike glycoprotein precursor | P59594.1 | Spike glycoprotein |
| 7 | 558456 | LYQDVNC | 597 | 603 | Spike glycoprotein precursor | P59594.1 | Spike glycoprotein |
| 8 | 558457 | LYQDVNCT | 597 | 604 | Spike glycoprotein precursor | P59594.1 | Spike glycoprotein |
| 9 | 18594 | GAGICASY | 653 | 660 | Spike glycoprotein precursor | P59594.1 | Spike glycoprotein |
| 10 | 22321 | GSFCTQLN | 739 | 746 | Spike glycoprotein precursor | P59594.1 | Spike glycoprotein |
| 11 | 16183 | FIEDLLFNKVTLADAGF | 799 | 815 | E2 glycoprotein precursor | NP_828851.1 | Spike glycoprotein |
| 12 | 4129 | ARDLICAQKFNGLTVLP | 828 | 844 | E2 glycoprotein precursor | NP_828851.1 | Spike glycoprotein |
| 13 | 18515 | GAALQIPFAMQMAYRFN | 873 | 889 | E2 glycoprotein precursor | NP_828851.1 | Spike glycoprotein |
| 14 | 3176 | AMQMAYRF | 881 | 888 | Spike glycoprotein precursor | P59594.1 | Spike glycoprotein |
| 15 | 10778 | DVVNQNAQALNTLVKQL | 932 | 948 | E2 glycoprotein precursor | NP_828851.1 | Spike glycoprotein |
| 16 | 50311 | QALNTLVKQLSSNFGAI | 939 | 955 | E2 glycoprotein precursor | NP_828851.1 | Spike glycoprotein |
| 17 | 33032 | KQLSSNFGAISSVLNDI | 946 | 962 | E2 glycoprotein precursor | NP_828851.1 | Spike glycoprotein |
| 18 | 11038 | EAEVQIDRLITGRLQSL | 970 | 986 | E2 glycoprotein precursor | NP_828851.1 | Spike glycoprotein |
| 19 | 54599 | RLITGRLQSLQTYVTQQ | 977 | 993 | E2 glycoprotein precursor | NP_828851.1 | Spike glycoprotein |
| 20 | 59425 | SLQTYVTQQLIRAAEIR | 985 | 1001 | E2 glycoprotein precursor | NP_828851.1 | Spike glycoprotein |
| 21 | 51379 | QLIRAAEIRASANLAAT | 993 | 1009 | E2 glycoprotein precursor | NP_828851.1 | Spike glycoprotein |
| 22 | 53202 | RASANLAATKMSECVLG | 1001 | 1017 | E2 glycoprotein precursor | NP_828851.1 | Spike glycoprotein |
| 23 | 462 | AATKMSECVLGQSKRVD | 1007 | 1023 | E2 glycoprotein precursor | NP_828851.1 | Spike glycoprotein |
| 24 | 69513 | VLGQSKRVDFCGKGYHL | 1015 | 1031 | E2 glycoprotein precursor | NP_828851.1 | Spike glycoprotein |
| 25 | 67220 | TVYDPLQPELDSFKEEL | 1118 | 1134 | E2 glycoprotein precursor | NP_828851.1 | Spike glycoprotein |
| 26 | 47341 | PELDSFKEELDKYFKNH | 1125 | 1141 | E2 glycoprotein precursor | NP_828851.1 | Spike glycoprotein |
| 27 | 10113 | DSFKEELDKYFKNHTSPDVDLGDISGINASVV | 1128 | 1159 | Spike glycoprotein precursor | P59594.1 | Spike glycoprotein |
| 28 | 11740 | EELDKYFKNHTSPDVDL | 1132 | 1148 | E2 glycoprotein precursor | NP_828851.1 | Spike glycoprotein |
| 29 | 9007 | DKYFKNHTSPDVDLGD | 1135 | 1150 | E2 glycoprotein precursor | NP_828851.1 | Spike glycoprotein |
| 30 | 32508 | KNHTSPDVDLGDISGIN | 1139 | 1155 | E2 glycoprotein precursor | NP_828851.1 | Spike glycoprotein |
| 31 | 60024 | SPDVDLGDISGINAS | 1143 | 1157 | Spike glycoprotein precursor | P59594.1 | Spike glycoprotein |
| 32 | 9094 | DLGDISGINASVVNIQK | 1147 | 1163 | E2 glycoprotein precursor | NP_828851.1 | Spike glycoprotein |
| 33 | 28512 | ISGINASVVNIQKEIDRLNE | 1151 | 1170 | E2 glycoprotein precursor | NP_828851.1 | Spike glycoprotein |
