## Supplemental Table 6 for "SARS-CoV2 spike protein displays biologically significant similarities with paramyxovirus surface proteins; a bioinformatics study"

### Supp Table 6- IEDB T cell epitopes with exact matches with substrings of SARS-CoV2 spike protein

| No | Epitope ID | Description | Starting Position | Ending Position | Antigen Name | Antigen Accession | Parent Protein |
|---|---|---|---|---|---|---|---|
| 1 | 70066 | VNFNFNGL | 525 | 532 | Spike glycoprotein precursor | P59594.1 | Spike glycoprotein |
| 2 | 100048 | GAALQIPFAMQMAYRF | 873 | 888 | S protein | ABF65836.1 | Spike glycoprotein |
| 3 | 50311 | QALNTLVKQLSSNFGAI | 939 | 955 | S protein | ABF65836.1 | Spike glycoprotein |
| 4 | 2801 | ALNTLVKQL | 940 | 948 | S protein | ABF65836.1 | Spike glycoprotein |
| 5 | 36724 | LITGRLQSL | 978 | 986 | Spike glycoprotein precursor | P59594.1 | Spike glycoprotein |
| 6 | 100428 | QLIRAAEIRASANLAATK | 993 | 1010 | S protein | ABF65836.1 | Spike glycoprotein |
| 7 | 71663 | VVFLHVTYV | 1042 | 1050 | Spike glycoprotein precursor | P59594.1 | Spike glycoprotein |
| 8 | 44814 | NLNESLIDL | 1174 | 1182 | S protein | ABF65836.1 | Spike glycoprotein |
| 9 | 16156 | FIAGLIAIV | 1202 | 1210 | Spike glycoprotein precursor | P59594.1 | Spike glycoprotein |
