## Supplemental Table 7 for "SARS-CoV2 spike protein displays biologically significant similarities with paramyxovirus surface proteins; a bioinformatics study"

### Supp Table 7- IEDB MHC binding epitopes with exact matches with substrings of SARS-CoV2 spike protein- Continued on next page

| No | Epitope ID | Description | Starting Position | Ending Position | Antigen Name | Antigen Accession | Parent Protein |
|---|---|---|---|---|---|---|---|
| 1 | 70718 | VRFPNITNL | 314 | 322 | Spike glycoprotein precursor | P59594.1 | Spike glycoprotein |
| 2 | 17341 | FPNITNLCPF | 316 | 325 | Spike glycoprotein precursor | P59594.1 | Spike glycoprotein |
| 3 | 7247 | CVADYSVLY | 348 | 356 | Spike glycoprotein precursor | P59594.1 | Spike glycoprotein |
| 4 | 23436 | GYQPYRVVVL | 490 | 499 | Spike glycoprotein precursor | P59594.1 | Spike glycoprotein |
| 5 | 51999 | QPYRVVVLSF | 492 | 501 | Spike glycoprotein precursor | P59594.1 | Spike glycoprotein |
| 6 | 50166 | PYRVVVLSF | 493 | 501 | Spike glycoprotein precursor | P59594.1 | Spike glycoprotein |
| 7 | 7289 | CVNFNFNGLTGTGVL | 524 | 538 | Spike glycoprotein precursor | P59594.1 | Spike glycoprotein |
| 8 | 47041 | PCSFGGVSVITPGTN | 575 | 589 | Spike glycoprotein precursor | P59594.1 | Spike glycoprotein |
| 9 | 69882 | VLYQDVNCT | 596 | 604 | Spike glycoprotein precursor | P59594.1 | Spike glycoprotein |
| 10 | 26198 | IGAGICASY | 652 | 660 | Spike glycoprotein precursor | P59594.1 | Spike glycoprotein |
| 11 | 37544 | LLLQYGSFC | 734 | 742 | Spike glycoprotein precursor | P59594.1 | Spike glycoprotein |
| 12 | 37724 | LLQYGSFCT | 735 | 743 | Spike glycoprotein precursor | P59594.1 | Spike glycoprotein |
| 13 | 22322 | GSFCTQLNR | 739 | 747 | Spike glycoprotein precursor | P59594.1 | Spike glycoprotein |
| 14 | 57792 | SFIEDLLFNK | 798 | 807 | Spike glycoprotein precursor | P59594.1 | Spike glycoprotein |
| 15 | 25754 | IEDLLFNKVTLADAG | 800 | 814 | Spike glycoprotein precursor | P59594.1 | Spike glycoprotein |
| 16 | 11384 | EDLLFNKVTLADAGF | 801 | 815 | Spike glycoprotein precursor | P59594.1 | Spike glycoprotein |
| 17 | 37289 | LLFNKVTLA | 803 | 811 | spike glycoprotein precursor | AEA10473.1 | Spike glycoprotein |
| 18 | 3982 | AQKFNGLTVLPPLLT | 834 | 848 | Spike glycoprotein precursor | P59594.1 | Spike glycoprotein |
| 19 | 23293 | GWTFGAGAALQIPFA | 867 | 881 | Spike glycoprotein precursor | P59594.1 | Spike glycoprotein |
| 20 | 18514 | GAALQIPFAMQMAYR | 873 | 887 | Spike glycoprotein precursor | P59594.1 | Spike glycoprotein |
| 21 | 38855 | LQIPFAMQM | 876 | 884 | spike glycoprotein | ABI96958.1 | Spike glycoprotein |
| 22 | 51112 | QIPFAMQMAYRFNGI | 877 | 891 | Spike glycoprotein precursor | P59594.1 | Spike glycoprotein |
| 23 | 65906 | TQNVLYENQK | 894 | 903 | Spike glycoprotein precursor | P59594.1 | Spike glycoprotein |
| 24 | 38831 | LQDVVNQNAQALNTL | 930 | 944 | Spike glycoprotein precursor | P59594.1 | Spike glycoprotein |
| 25 | 3939 | AQALNTLVK | 938 | 946 | Spike glycoprotein precursor | P59594.1 | Spike glycoprotein |
| 26 | 2801 | ALNTLVKQL | 940 | 948 | Spike glycoprotein precursor | P59594.1 | Spike glycoprotein |
| 27 | 38353 | LNTLVKQLSSNFGAI | 941 | 955 | Spike glycoprotein precursor | P59594.1 | Spike glycoprotein |
| 28 | 923559 | NFGAISSVL | 951 | 959 | Spike glycoprotein precursor | P59594.1 | Spike glycoprotein |
| 29 | 54507 | RLDKVEAEV | 965 | 973 | Spike glycoprotein precursor | P59594.1 | Spike glycoprotein |
| 30 | 1220 | AEVQIDRLI | 971 | 979 | Spike glycoprotein precursor | P59594.1 | Spike glycoprotein |
| 31 | 1221 | AEVQIDRLIT | 971 | 980 | Spike glycoprotein precursor | P59594.1 | Spike glycoprotein |
| 32 | 70575 | VQIDRLITGR | 973 | 982 | Spike glycoprotein precursor | P59594.1 | Spike glycoprotein |
| 33 | 25662 | IDRLITGRLQSLQTY | 975 | 989 | Spike glycoprotein precursor | P59594.1 | Spike glycoprotein |
| Continued | | | | | | | |

### Supp Table 7- Continued

| No | Epitope ID | Description | Starting Position | Ending Position | Antigen Name | Antigen Accession | Parent Protein |
|---|---|---|---|---|---|---|---|
| 34 | 36724 | LITGRLQSL | 978 | 986 | Spike glycoprotein precursor | P59594.1 | Spike glycoprotein |
| 35 | 63951 | TGRLQSLQTYVTQQL | 980 | 994 | Spike glycoprotein precursor | P59594.1 | Spike glycoprotein |
| 36 | 22144 | GRLQSLQTY | 981 | 989 | Spike glycoprotein precursor | P59594.1 | Spike glycoprotein |
| 37 | 54725 | RLQSLQTYV | 982 | 990 | spike glycoprotein precursor | AEA10473.1 | Spike glycoprotein |
| 38 | 38990 | LQSLQTYVTQQLIRA | 983 | 997 | Spike glycoprotein precursor | P59594.1 | Spike glycoprotein |
| 39 | 39003 | LQTYVTQQLIRAAEI | 986 | 1000 | Spike glycoprotein precursor | P59594.1 | Spike glycoprotein |
| 40 | 52672 | QTYVTQQLIRAAEIR | 987 | 1001 | Spike glycoprotein precursor | P59594.1 | Spike glycoprotein |
| 41 | 52057 | QQLIRAAEIRASANL | 992 | 1006 | Spike glycoprotein precursor | P59594.1 | Spike glycoprotein |
| 42 | 999 | AEIRASANLA | 998 | 1007 | Spike glycoprotein precursor | P59594.1 | Spike glycoprotein |
| 43 | 4321 | ASANLAATK | 1002 | 1010 | Spike glycoprotein precursor | P59594.1 | Spike glycoprotein |
| 44 | 56252 | RVDFCGKGY | 1021 | 1029 | Spike glycoprotein precursor | P59594.1 | Spike glycoprotein |
| 45 | 3589 | APHGVVFLHV | 1038 | 1047 | Spike glycoprotein precursor | P59594.1 | Spike glycoprotein |
| 46 | 23200 | GVVFLHVTY | 1041 | 1049 | Spike glycoprotein precursor | P59594.1 | Spike glycoprotein |
| 47 | 71663 | VVFLHVTYV | 1042 | 1050 | Spike glycoprotein precursor | P59594.1 | Spike glycoprotein |
| 48 | 71996 | VYDPLQPEL | 1119 | 1127 | Spike glycoprotein precursor | P59594.1 | Spike glycoprotein |
| 49 | 10112 | DSFKEELDKY | 1128 | 1137 | Spike glycoprotein precursor | P59594.1 | Spike glycoprotein |
| 50 | 35205 | LDKYFKNHTSPDVDL | 1134 | 1148 | Spike glycoprotein precursor | P59594.1 | Spike glycoprotein |
| 51 | 9006 | DKYFKNHTSPDVDLG | 1135 | 1149 | Spike glycoprotein precursor | P59594.1 | Spike glycoprotein |
| 52 | 36075 | LGDISGINASVVNIQ | 1148 | 1162 | Spike glycoprotein precursor | P59594.1 | Spike glycoprotein |
| 53 | 28511 | ISGINASVVNIQKEI | 1151 | 1165 | Spike glycoprotein precursor | P59594.1 | Spike glycoprotein |
| 54 | 44814 | NLNESLIDL | 1174 | 1182 | Spike glycoprotein precursor | P59594.1 | Spike glycoprotein |
| 55 | 59161 | SLIDLQELGK | 1178 | 1187 | Spike glycoprotein precursor | P59594.1 | Spike glycoprotein |
| 56 | 36481 | LIDLQELGKY | 1179 | 1188 | Spike glycoprotein precursor | P59594.1 | Spike glycoprotein |
| 57 | 50641 | QELGKYEQYI | 1183 | 1192 | Spike glycoprotein precursor | P59594.1 | Spike glycoprotein |
| 58 | 73751 | YEQYIKWPWY | 1188 | 1197 | Spike glycoprotein precursor | P59594.1 | Spike glycoprotein |
| 59 | 72717 | WLGFIAGLIAIVMVT | 1199 | 1213 | Spike glycoprotein precursor | P59594.1 | Spike glycoprotein |
| 60 | 36103 | LGFIAGLIAIVMVTI | 1200 | 1214 | Spike glycoprotein precursor | P59594.1 | Spike glycoprotein |
| 61 | 16156 | FIAGLIAIV | 1202 | 1210 | Spike glycoprotein precursor | P59594.1 | Spike glycoprotein |
| 62 | 20907 | GLIAIVMVTI | 1205 | 1214 | spike glycoprotein precursor | ADC35483.1 | Spike glycoprotein |
| 63 | 6668 | CMTSCCSCLK | 1218 | 1227 | Spike glycoprotein precursor | P59594.1 | Spike glycoprotein |
| 64 | 42873 | MTSCCSCLK | 1219 | 1227 | Spike glycoprotein precursor | P59594.1 | Spike glycoprotein |
| 65 | 57592 | SEPVLKGVKL | 1243 | 1252 | Spike glycoprotein precursor | P59594.1 | Spike glycoprotein |
