## Supplemental Table 8 for "SARS-CoV2 spike protein displays biologically significant similarities with paramyxovirus surface proteins; a bioinformatics study"

### Supp Table 8- representative ssRNA viruses used for phylogenetic analyses

| Sense | Family | Virus |
|---|---|---|
| Negative | Orthomyxoviridae | Influenza A virus |
| Negative | Orthomyxoviridae | Influenza B virus |
| Negative | Orthomyxoviridae | Influenza C virus |
| Negative | Orthomyxoviridae | Influenza D virus |
| Negative | Orthomyxoviridae | Salmon Isavirus |
| Negative | Paramyxoviridae | Measles morbillivirus |
| Negative | Paramyxoviridae | Rinderpest morbillivirus |
| Negative | Paramyxoviridae | Mumps rubulavirus |
| Negative | Paramyxoviridae | Parainfluenza virus 1 (Human respirovirus 1) |
| Negative | Paramyxoviridae | Parainfluenza virus 2 (Human rubulavirus 2) |
| Negative | Paramyxoviridae | Parainfluenza virus 3 (Human respirovirus 3) |
| Negative | Paramyxoviridae | Parainfluenza virus 5 (Mammalian rubulavirus 5) |
| Negative | Filoviridae | Zaire ebolavirus |
| Negative | Filoviridae | Marburg marburgvirus |
| Negative | Filoviridae | Lloviu cuevavirus |
| Negative | Filoviridae | Mengla dianlovirus |
| Negative | Rhabdoviridae | Rabies lyssavirus |
| Negative | Rhabdoviridae | Vesicular stomatitis Indiana virus |
| Negative | Rhabdoviridae | European bat lyssa virus 1 |
| Negative | Rhabdoviridae | Bas-Congo tibrovirus |
| Negative | Rhabdoviridae | Viral hemorrhagic septicemia virus |

| Sense | Family | Virus |
|---|---|---|
| Positive | Coronaviridae | SARS-CoV2 |
| Positive | Coronaviridae | SARS -CoV |
| Positive | Coronaviridae | MERS-related coronavirus |
| Positive | Coronaviridae | Human coronavirus OC43 |
| Positive | Coronaviridae | Human coronavirus HKU1 |
| Positive | Coronaviridae | Porcine epidemic diarrhea virus |
| Positive | Coronaviridae | Murine hepatitis virus |
| Positive | Picornaviridae | Enterovirus, EV-71 |
| Positive | Picornaviridae | Enterovirus, Coxackie virus |
| Positive | Picornaviridae | Enterovirus, poliovirus |
| Positive | Picornaviridae | Enterovirus, Rhinovirus A |
| Positive | Picornaviridae | Enterovirus B |
| Positive | Picornaviridae | Hepatovirus A, hep A virus |
| Positive | Picornaviridae | Aphthovirus (Equine rhinitis A virus) |
| Positive | Flaviviridae | Hepatitis C virus |
| Positive | Flaviviridae | Zika virus |
| Positive | Flaviviridae | Dengue virus 2 |
| Positive | Flaviviridae | West Nile virus |
| Positive | Flaviviridae | Bovine viral diarrhea virus 1 |
| Positive | Togaviridae | Semliki Forest virus |
| Positive | Togaviridae | Sindbis virus |
| Positive | Togaviridae | Eastern equine encephalitis virus |
| Positive | Togaviridae | Western equine encephalitis virus |
| Positive | Togaviridae | Rubella virus |
| Positive | Caliciviridae | Norovirus GI |
| Positive | Caliciviridae | Feline calicivirus |
| Positive | Caliciviridae | Sapovirus |
| Positive | Caliciviridae | Primate norovirus |
| Positive | Caliciviridae | Canine vesivirus |
| Positive | Caliciviridae | Norwalk like virus |
